# Leveraging lung tissue transcriptome to uncover candidate causal genes in COPD genetic associations

**DOI:** 10.1101/193938

**Authors:** Ma’en Obeidat, Maxime Lamontagne, Jean-Christophe Bérubé, Michael H Cho, Brian D. Hobbs, Kim de Jong, H. Marike Boezen, the International COPD Genetics Consortium, David Nickle, Ke Hao, Wim Timens, Maarten van den Berge, Philippe Joubert, Michel Laviolette, Don D Sin, Peter D Paré, Yohan Bossé

## Abstract

We collated 129 non-overlapping risk loci for chronic obstructive pulmonary disease (COPD) from the GWAS literature. Using recent and complementary integrative genomics approaches, combining GWAS and lung eQTL results, we identified 12 novel COPD loci and corresponding causal genes. In addition, we mapped candidate causal genes for 60 out of the 129 GWAS-nominated loci as well as for four sub-genome-wide significant COPD risk loci derived from the largest GWAS on COPD. Mapping causal genes in lung tissue represents an important contribution on the genetics of COPD, enriches our biological interpretation of GWAS findings, and brings us closer to clinical translation of genetic associations.

## INTRODUCTION

Chronic obstructive pulmonary disease (COPD) is among the leading causes of hospitalization in industrialized countries and is the third leading cause of death worldwide.^1^ It was recently estimated that the absolute number of COPD cases in developed countries will increase by more than 150% from 2010 to 2030.^2^ The lack of understanding of the molecular mechanisms underlying the pathogenesis of COPD has hampered efforts to develop new biomarkers and effective therapies.

Cigarette smoking is the main modifiable environmental risk factor for COPD. However, only 20-25% of smokers develop clinically significant airflow obstruction.^3^ There is strong evidence for genetic contribution to COPD. Candidate gene and genome-wide association studies (GWAS) have identified many genetic variants associated with COPD and its related phenotypes.^4-9^ The latest GWAS was performed by the International COPD Genetics Consortium (ICGC) which included 15,256 cases and 47,936 controls, and replication of significant signals in additional 9,498 cases and 9,748 controls.^10^ The combined meta-analysis identified 22 COPD susceptibility loci at genome-wide significance. However, it is likely that many additional loci contributing to COPD pathogenesis were missed because of the stringent statistical threshold typically used in GWAS.

Biological interpretation of genetic association results remains a major challenge. Most GWAS associated variants have regulatory function and are associated with changes in gene expression.^11,12^ The mapping of expression quantitative trait loci (eQTL) in disease-relevant tissues has been successfully used to identify the causal genes underpinning GWAS-nominated loci.^13^ Using lung eQTL derived from 1,038 subjects,^14^ we have previously identified the likely causal genes within COPD susceptibility loci.^15-18^ Recently developed methods allow more advanced integration of GWAS and eQTL results to colocalize GWAS and eQTL signals^19,20^ as well as to perform transcriptome-wide association study (TWAS)^21^ to identify the causal genes and functional SNPs underlying biological traits and diseases. In this study, we used three complementary methods to integrate the ICGC COPD GWAS results^10^ and previously published COPD loci with lung tissue eQTLs^14^ to identify the most likely causal COPD genes.

## RESULTS

### Overall study design

The study design is shown in **Fig. 1**. The ICGC COPD GWAS and lung eQTL datasets were integrated using three methods: TWAS, colocalisation, and Mendelian randomization-based (SMR) approach. These methods were first applied at the genome-wide level to identify new COPD loci/genes. Results were then further evaluated in risk loci derived from published literature of GWAS on COPD and related phenotypes as well as to liberal COPD risk loci (P_GWAS_<1x10^−5^, but sub-genome-wide significant) from ICGC that were non-overlapping with published GWAS loci. Finally, direct eQTL evaluation of GWAS SNPs (eSNP) was also assessed for literature-based and liberal COPD loci.

**Figure 1.**
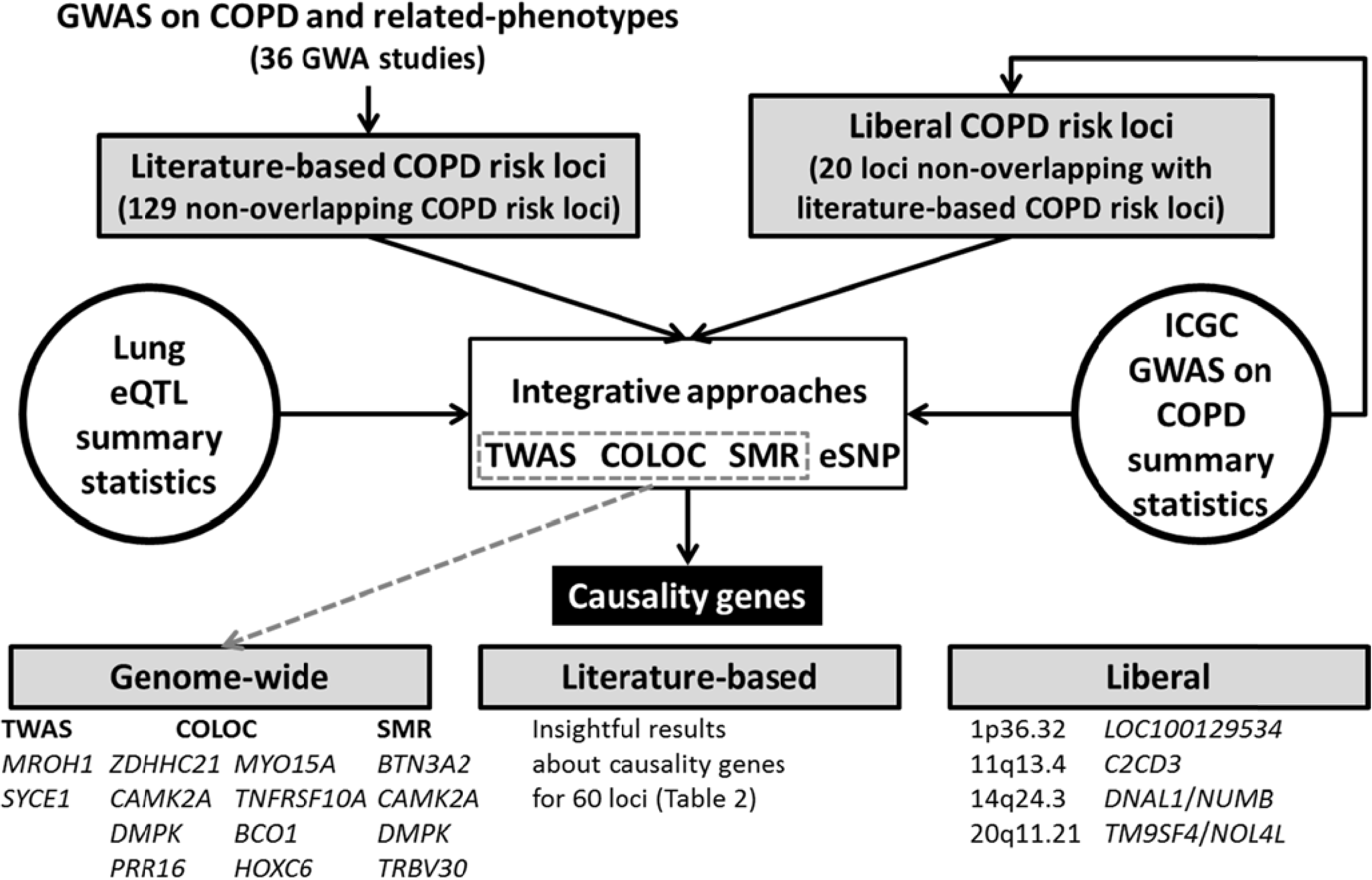
Overview of the study design.

### Genome-wide integrative approaches

#### TWAS

A Manhattan plot showing transcriptome-wide associations in lung tissue with COPD is shown in **Fig. 2a**. The 11 gene-COPD associations (corresponding to 16 probe sets) that reached genome-wide significance (P_TWAS_<0.0001, **Supplementary Table 1**) are shown. Of these, 12 probe sets resided in literature-based GWAS loci including four on 6p24.3, two on 5q32, two on 16p11.2, and two on 19q13.2. In contrast, *MROH1* on 8q24.3 and *SYCE1* on 10q26.3 represented novel candidate causal genes for COPD (**Table 1**).

**Table 1.** Twelve novel COPD susceptibility genes/loci.

| **Gene** | **Locus** | **P<sub>TWAS</sub>** | **PP4<sub>COLOC</sub>** | **P<sub>SMR</sub>** | **P<sub>eQTL</sub> (SNP)<sup>a</sup>** | **Gene function<sup>b</sup>** |
| --- | --- | --- | --- | --- | --- | --- |
| *MROH1* | 8q24.3 | **9.65E-09** | 0.03 | - | **2.868E-60**<br>(rs113954825) | Lysosomal regulator |
| *SYCE1* | 10q26.3 | **4.93E-05** | 0.05 | - | 1.286E-08<br>(rs146332556) | Meiosis, cancer |
| *ZDHHC21* | 9p22.3 | 0.886 | **0.95** | 0.0034 | **2.988E-25**<br>(rs10756585) | Endothelial barrier integrity |
| *CAMK2A* | 5q32 | 0.260 | **0.91** | **0.0006** | **6.611E-14**<br>(rs930212) | Calcium signaling |
| *DMPK* | 19q13.32 | 0.087 | **0.86** | **0.0009** | **2.055E-23**<br>(rs7253302) | Antioxidant, development |
| *PRR16* | 5q23.1 | 0.165 | **0.82** | - | 7.466E-07<br>(rs10053752) | Regulator of cell size |
| *MYO15A* | 17p11.2 | 0.277 | **0.81** | 0.0050 | **1.98E-12**<br>(rs9916193) | Actin-organization in inner ear hair cells |
| *TNFRSF10A* | 8p21.3 | 0.404 | **0.81** | 0.0033 | **8.191E-32**<br>(rs13278062) | Apoptosis |
| *BCO1* | 16q23.2 | 0.962 | **0.77** | - | 5.72E-07<br>(rs72823838) | Alveolar development and repair |
| *HOXC6* | 12q13.13 | 0.891 | **0.75** | - | 7.881E-07<br>(rs746423) | Development and cancer |
| *BTN3A2* | 6p22.2 | 0.011 | 0.72 | **0.0003** | **6.10E-240**<br>(rs9366653) | Immune system, anti-tumor |
| *TRBV30* | 7q34 | 0.262 | 0.74 | **0.0009** | **3.211E-39**<br>(rs17267) | Immune response |

P<sub>TWAS</sub><0.0001, PP4<0.75, P<sub>SMR</sub><0.001, and P<sub>eQTL</sub><1x10<sup>-8</sup> are in bold.
<sup>a</sup>Top eQTL for the probe set.
<sup>b</sup>See **Supplementary Table 10** for more details.

**Figure 2.**
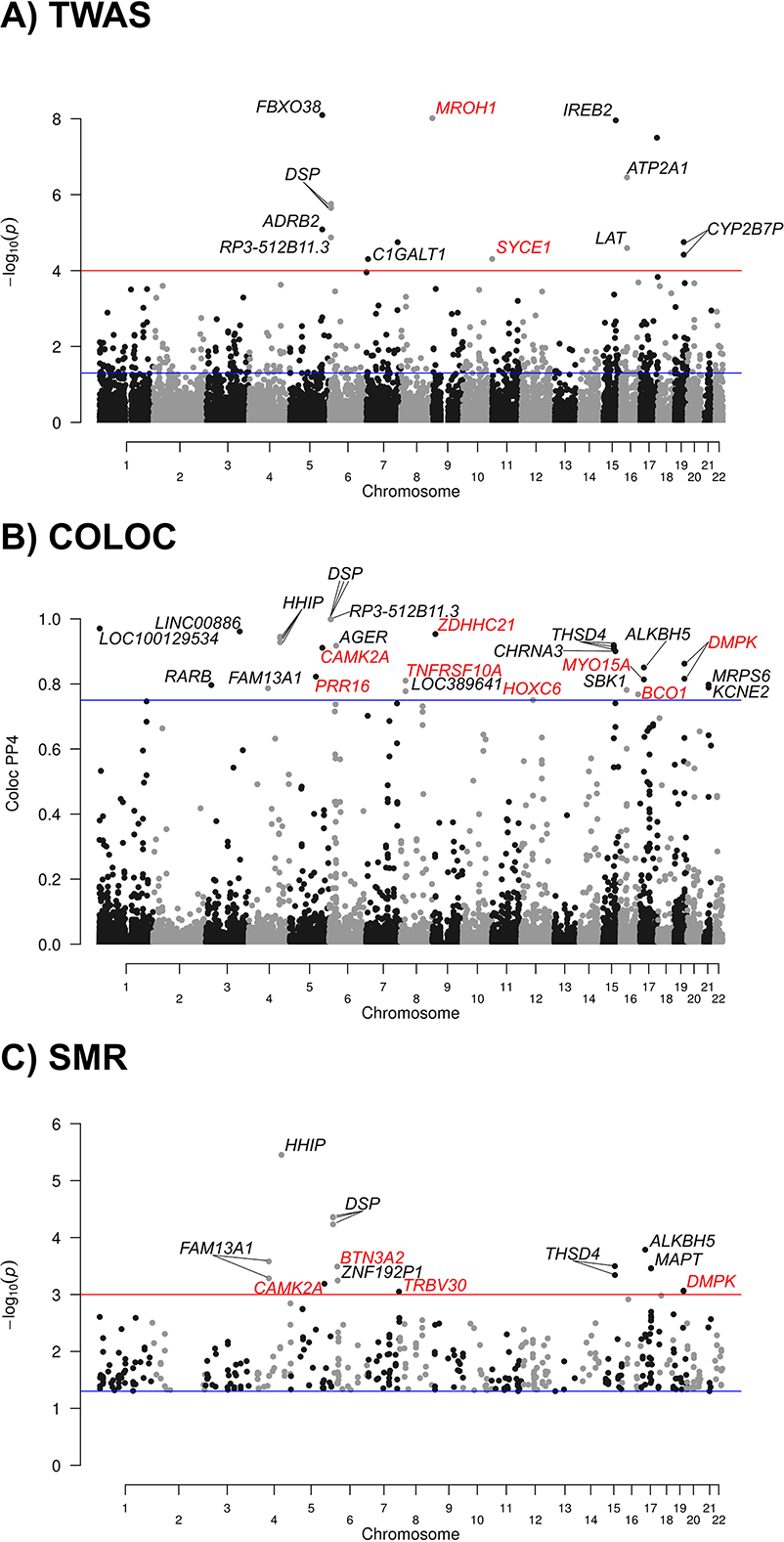
Manhattan plots of TWAS, colocalisation, and SMR. **A**) TWAS results showing p values on the –log10 scale on the y-axis for gene expression-COPD associations. The blue horizontal line represents P_TWAS_ of 0.05. The red horizontal line represents the genome-wide significant threshold used in this study (P_TWAS_<0.0001). Annotations for significant probe sets are indicated. **B**) Colocalisation results showing posterior probability of shared signals (PP4) on the y-axis. The blue horizontal line represents a standard colocalisation threshold of 75%. Probe sets above this threshold are annotated. **C**) SMR results showing p values on the –log10 scale on the y-axis. By default, SMR only considered probe sets with at least one *cis*-eQTL<5.0x10^−8^. The blue horizontal line represents P_SMR_ of 0.05. The red horizontal line represents the significant threshold used in this study (P_SMR_<0.001). Probe sets showing heterogeneity (P_HEIDI_ < 0.05) were excluded. Probe sets with P_SMR_>0.05 were also removed as no heterogeneity (HEIDI) tests were performed. Gene annotations for probe sets with P_SMR_<0.001 are indicated. In the three Manhattan plots, gene names in red are new gene/loci discovered in this study.

#### Colocalisation

A Manhattan plot showing colocalisation results is shown in **Fig. 2b**. Posterior probability of shared signals (PP4) >75% are observed at 18 loci (**Supplementary Table 2**). Ten of them colocalised in literature-based and liberal COPD risk loci. The eight others represent novel candidate causal genes for COPD including *ZDHHC21* on 9p22.3, *CAMK2A* on 5q32, *DMPK* on 19q13.32, *PRR16* on 5q23.1, *MYO15A* on 17p11.2, *TNFRSF10A* on 8p21.3, *BCO1* on 16q23.2, and *HOXC6* on 12q13.13 (**Table 1**).

**Table 2.**
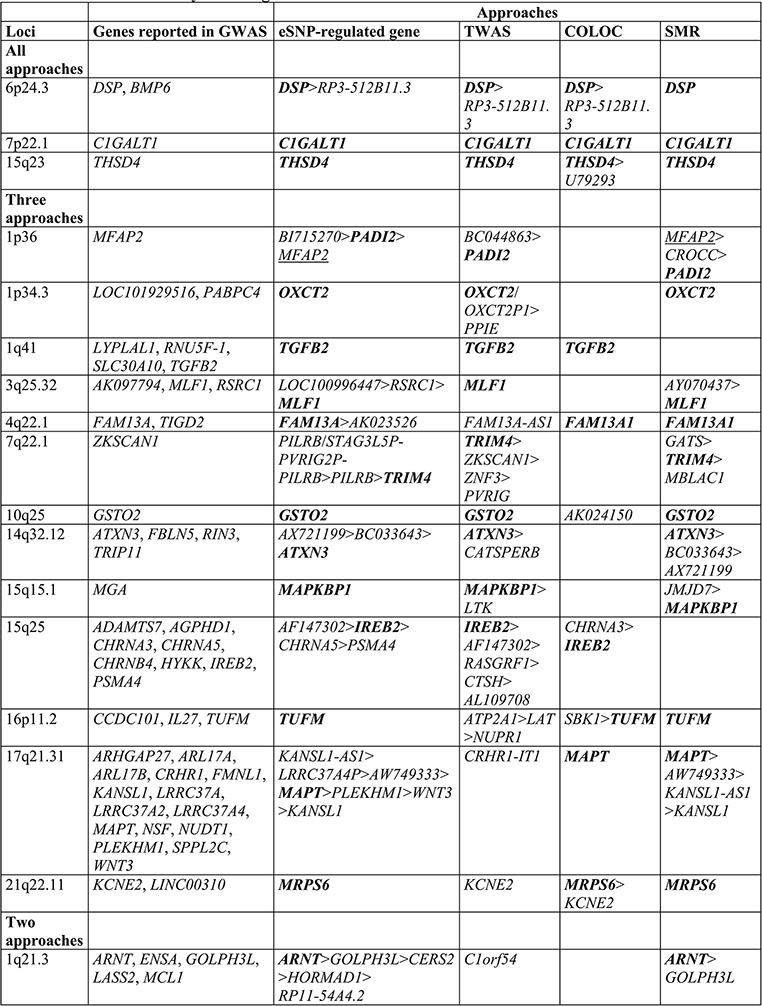
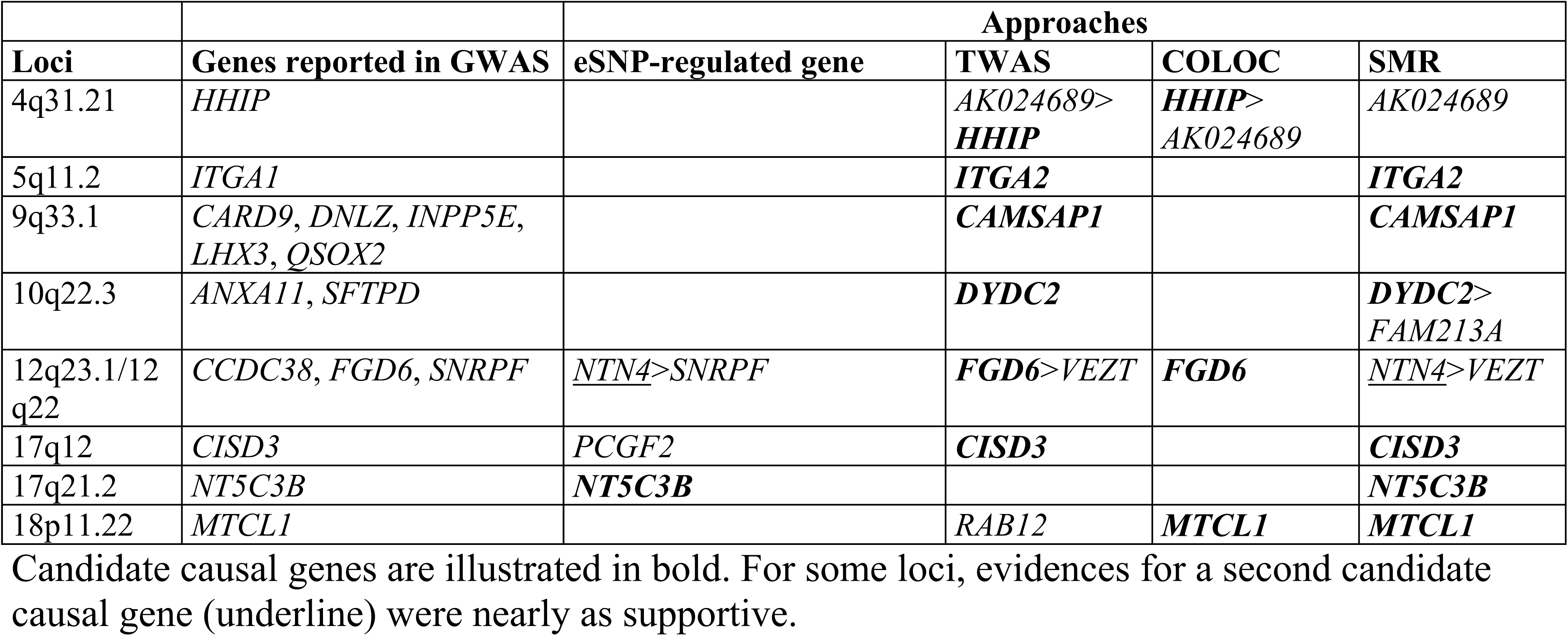
The most likely causal genes in literature-based COPD risk loci.

#### SMR

**Fig. 2c** and **Supplementary Table 3** show significant genes identified using the SMR method. Excluding the known COPD risk loci, the four genome-wide significant SMR genes (P_SMR_=0.001) were *BTN3A2* on 6p22.2, *CAMK2A* on 5q32, *DMPK* on 19q13.32, and *TRBV30* on 7q34. The four genes were also supported by colocalisation (PP4>0.72) (**Table 1**).

### Literature-based COPD risk loci

#### Lung eQTL

The results of 36 GWAS on COPD and related phenotypes are summarized in **Supplementary Table 4**. Risk loci were extracted from publications as well as key genes and SNPs reported/discussed by the authors. For the top GWAS SNP at each reported locus, we tested its effect on lung gene expression in *cis* (1 Mb distance on either side of the SNP). All eSNP-regulated genes with P_eQTL_<1x10^−8^ are reported in **Supplementary Table 5**. Top eSNP-regulated genes (ones with the lowest lung eQTL p value) included *KANSL1-AS1* and *MAPT* on chromosome 17q21.31, *DSP* on 6p24.3, *HSD17B12* on 11p11.2, *PSORS1C3* on 6p21.33, *ARNT* on 1q21.3, *HLA-DQB2* on 6p21.32, and *IREB2* on 15q25 (**Supplementary Fig. 1**). All eSNP-regulated genes with a nominal P_eQTL_<0.05 by loci and studies are reported in **Supplementary Table 4**.

#### Combining results from different approaches

GWAS results were arranged in 129 non-overlapping COPD risk loci (**Supplementary Table 6**). For each locus, GWAS summary statistics from ICGC and the lung eQTL study were integrated to find the most likely causal gene(s) using TWAS, colocalisation, SMR, and eSNP-regulated gene approaches. For all loci, **Supplementary Table 6** provides the boundaries of each locus, lead GWAS SNP in ICGC, top lung eQTL SNP, TWAS genes (P_TWAS_<0.05), colocalising genes (PP4>60%), and SMR genes (P_SMR_<0.05). **Table 2** summarizes loci for which insightful results were obtained about the candidate causal gene and that were consistent for at least two integrative approaches. **Supplementary Table 7** presents results for the 129 loci. The most consistent candidate causal genes identified in this study were *DSP* on 6p24.3, *C1GALT1* on 7p22.1, and *THSD4* on 15q23. For these three loci, TWAS, colocalisation, SMR, and direct eSNP assessment consistently converged on these three potential causal genes (**Fig. 3** and **Supplementary Fig. 2**). For 13 additional loci, the same candidate causal gene was identified by three approaches (**Supplementary Fig. 3**). Direct lung eQTL assessment of GWAS SNP, TWAS, and SMR supported *PADI2* on 1p36 (**Supplementary Fig. 3a**), *OXCT2* on 1p34.3 (**Supplementary Fig. 3b**), *MLF1* on 3q25.32 (**Supplementary Fig. 3d**), *TRIM4* on 7q22.1 (**Supplementary Fig. 3f**), *GSTO2* on 10q25 (**Supplementary Fig. 3g**), *ATXN3* on 14q32.12 (**Supplementary Fig. 3h**), and *MAPKBP1* on 15q15.1 (**Supplementary Fig. 3i**). Direct lung eQTL assessment of GWAS SNP, TWAS, and colocalisation supported *TGFB2* on 1q41 (**Supplementary Fig. 3c**) and *IREB2* on 15q25 (**Supplementary Fig. 3j**). Direct lung eQTL assessment of GWAS SNP, colocalisation and SMR supported *FAM13A1* on 4q22.1 (**Supplementary Fig. 3e**), *TUFM* on 16q11.2 (**Supplementary Fig. 3k**), *MAPT* on 17q21.31 (**Supplementary Fig. 3l**), and *MRPS6* on 21q22.11 (**Supplementary Fig. 3m**). Candidate causal genes supported by at least two approaches were found at nine other loci: *ARNT* on 1q21.3, *HHIP* on 4q31.21, *ITGA2* on 5q11.2, *CAMSAP1* on 9q33.1, *DYDC2* on 10q22.3, *FGD6* on 12q23.1-q22, *CISD3* on 17q12, *NT5C3B* on 17q21.2, and *MTCL1* on 18p11.22 (**Table 2**). Possible causal genes supported by a single approach were found at 35 loci, including 8 lung eSNP-regulated genes, 17 TWAS genes, 1 colocalisation gene, and 9 SMR genes (**Table 2**). Overall, insightful results about the causal genes were provided for 60 loci. For 23 of these (38%), the target gene was different from that reported in the GWAS (**Supplementary Fig. 4**). For 18 loci (30%), the suspected gene from the GWAS was confirmed. Finally, for 19 loci (32%), the investigation refined the search to a single gene among the list of genes suspected by the GWAS.

**Figure 3.**
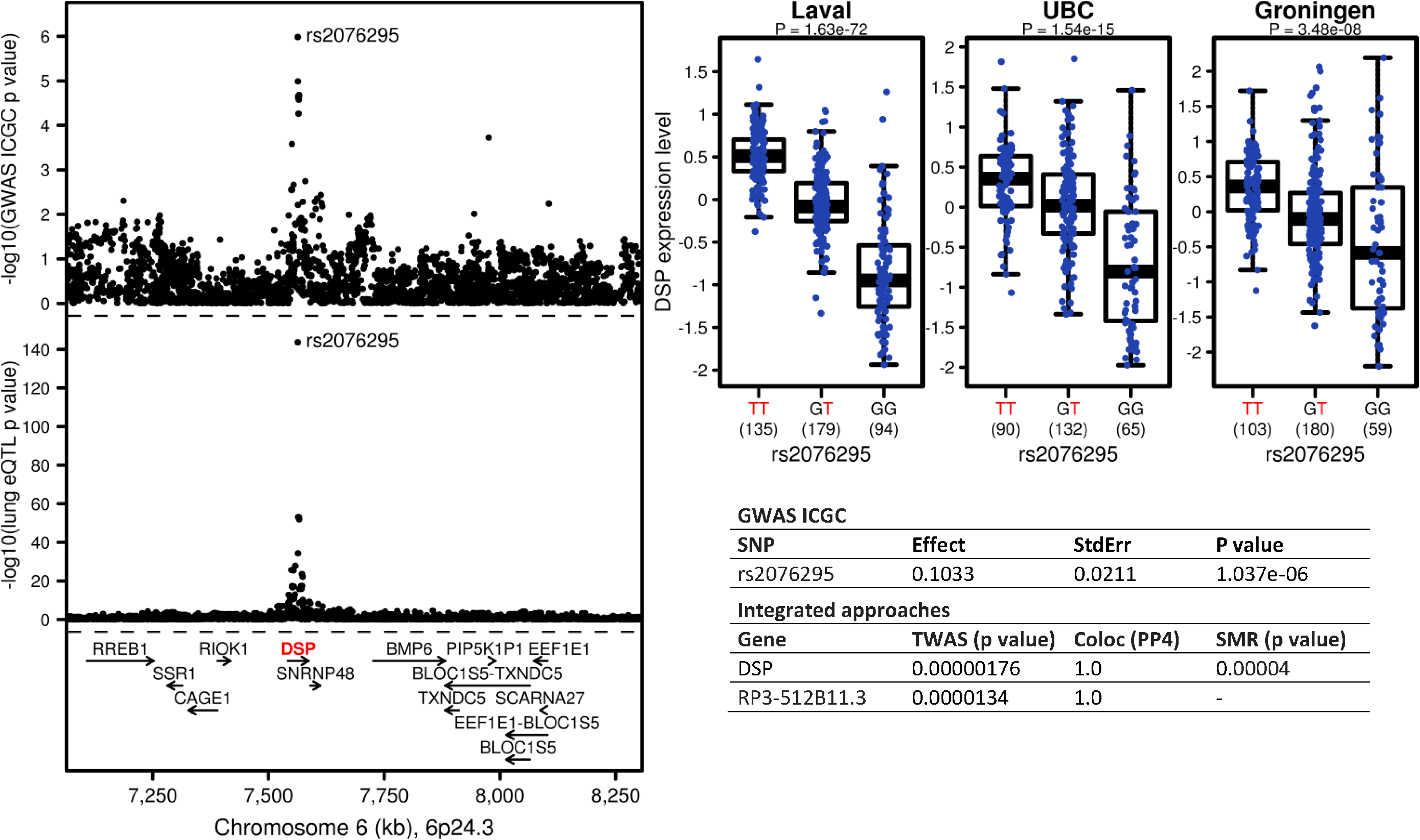
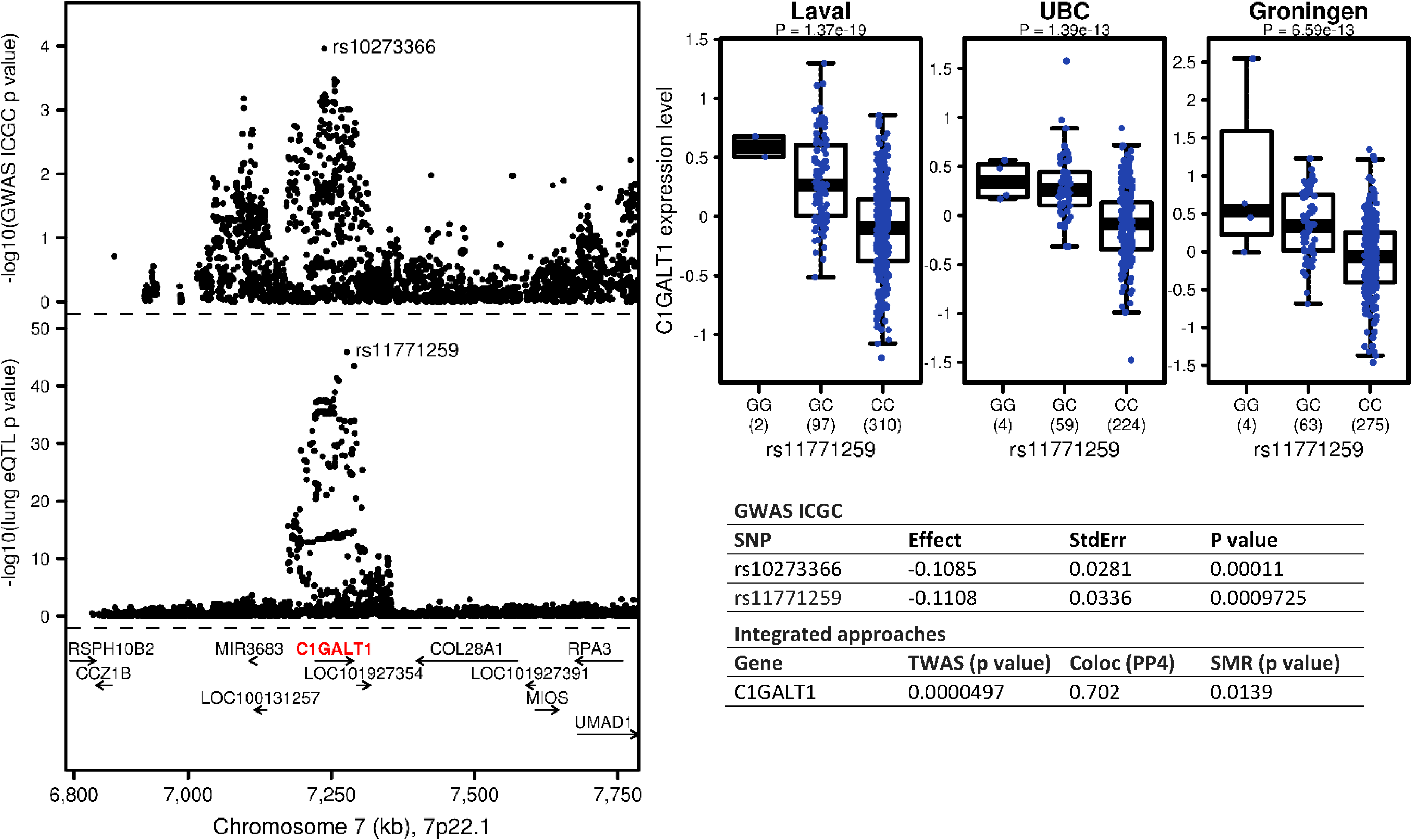
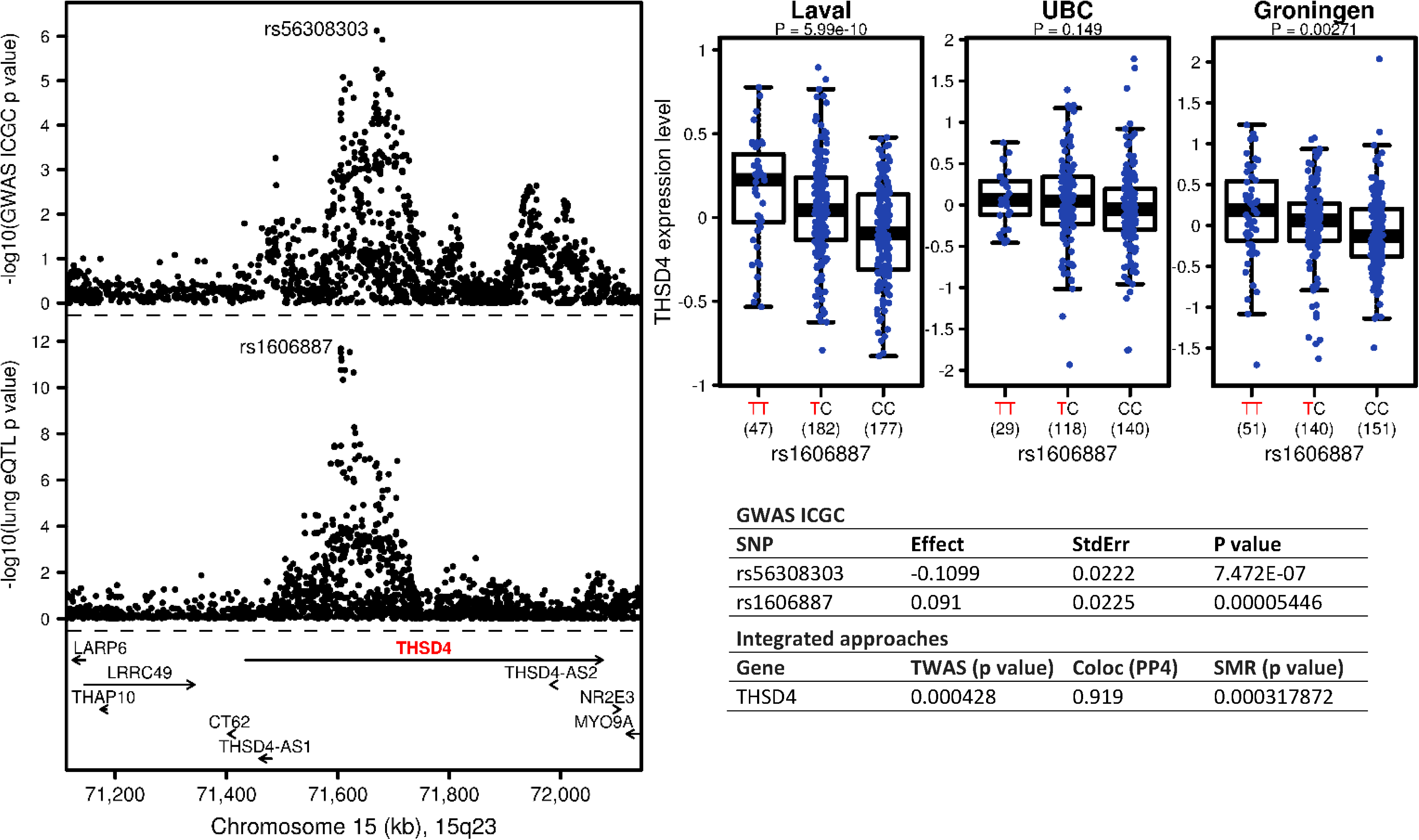
The most consistent candidate causal genes identified in this study. **A)*DSP* on chromosome 6p24.3, B)*C1GALT1* on chromosome 7p22.1, and C)*THSD4* on 15q23**. The upper left panel shows the genetic associations with COPD in ICGC. The bottom left panel shows the lung eQTL statistics for the target genes (*DSP*, *C1GALT1*, or *THSD4*). The boundaries of the loci are defined in **Supplementary Table 6** and the location of genes in this locus is illustrated at the bottom. The upper right panel shows boxplots of gene expression levels in the lung according to genotype groups for Laval, University of British Columbia (UBC) and Groningen samples. The y-axis shows the mRNA expression levels. The x-axis represents the three genotype groups for the SNP most strongly associated with mRNA expression of the target genes with the number of individuals in parenthesis. The risk allele identified in the ICGC GWAS is shown in red. Box boundaries, whiskers and centre mark in boxplots represent the first and third quartiles, the most extreme data point which is no more than 1.5 times the interquartile range, and the median, respectively. The table shows the top GWAS SNPs in ICGC and then the most likely causal gene(s) based on TWAS, colocalisation, and SMR combining summary statistics at these loci from the ICGC GWAS and the lung eQTL study. The linkage disequilibrium plots of selected SNPs on 6p24.3, 7p22.1, and 15q23 loci are provided in **Supplementary Fig. 2**.

**Figure 4.**
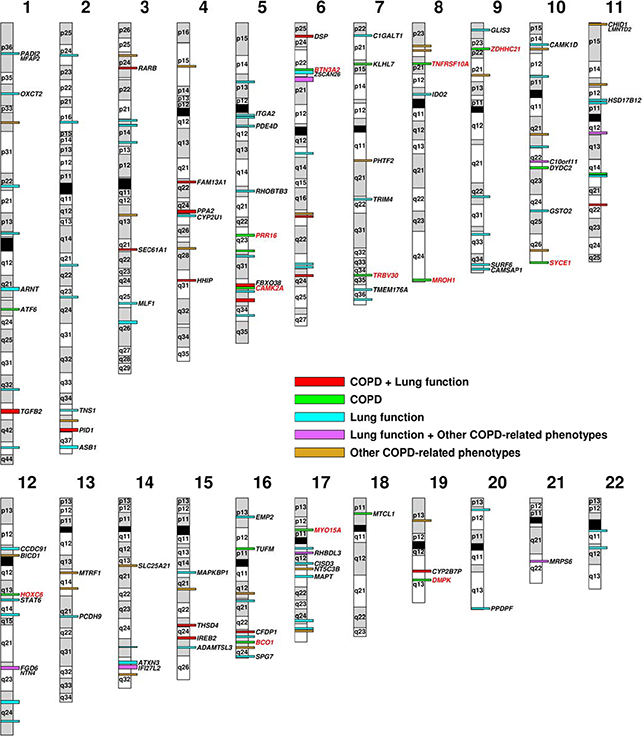
The COPD Gene Map. The map is an ideogram of the 22 autosomal human chromosomes and shows the location of candidate causal genes identified in this study. The map includes 12 genes in red located in new COPD susceptibility loci. The 129 non-overlapping COPD risk loci derived from published GWAS on COPD, lung function, and related phenotypes are also illustrated and are colored coded based on phenotypes (see color key). For 60 of these loci, the candidate causal genes identified in this study are indicated. The boundaries of each locus are provided in **Supplementary Table 6**. The alternating grey and white colors on the chromosomes have been used to distinguish cytogenic bands from the adjacent ones and do not correspond to the band colors observed on giamsa-stained chromosomes. Information to construct the ideogram was obtained from the UCSC Genome Browser (hg19).

The twelve novel COPD susceptibility genes/loci identified in this study are mapped in **Fig. 4**. In addition, **Fig. 4** illustrates the 129 non-overlapping COPD risk loci derived from GWAS as well as the corresponding candidate causal genes for 60 of them revealed in this study.

### Liberal COPD risk loci

Sub-genome-wide significant loci in the ICGC GWAS (5x10^−8^>P_GWAS_<1x10^−5^) were integrated with lung eQTL. A total of 40 loci were identified with at least one SNP associated with COPD at P_GWAS_<1x10^−5^ (**Supplementary Table 8**). Twenty loci were non-overlapping with literature-based COPD risk loci. Lung eSNP-regulated genes in these loci are shown in **Supplementary Table 9**. The most significant based on eQTL p value included the pseudogene *LOC100129534* on chromosome 1p36.32, *TM9SF4* on 20q11.21, and *DNAL1* on 14q24.3 (**Supplementary Fig. 5**). TWAS, colocalisation, and SMR results were further explored for liberal COPD risk loci. Three TWAS genes were identified including *C2CD3* on 11q13.4 (P_TWAS_=0.007), *NUMB* on 14q24.3 (P_TWAS_=0.021), and *NOL4L* on 20q11.21 (P_TWAS_=0.018). The eSNP-regulated pseudogene described above on 1p36.32 (*LOC100129534*) was also confirmed by colocalisation (PP4=0.97) and SMR (P_SMR_=0.0025).

## DISCUSSION

This is a comprehensive study to investigate the regulatory mechanisms in lung tissue underlying GWAS loci for COPD and its related phenotypes. Genome-wide integration of the largest GWAS on COPD with the largest lung eQTL study revealed 12 novel loci/genes causally related to COPD. This study also summarized 129 susceptibility loci from published GWAS on COPD and related phenotypes, and provided insightful results about the most likely causal genes for 60 (47%) of them. Liberal COPD susceptibility loci from the recent ICGC GWAS were also reported including 20 non-overlapping loci with literature-based GWAS. Plausible causality genes were revealed for four (20%) of these sub-genome-wide significant COPD loci. Finally, while the results from each method were slightly different, TWAS, SMR, COLOC, and direct eSNP assessment converged on three genes: *DSP* on 6p24.3, *C1GALT1* on 7p22.1, and *THSD4* on 15q23.

At the genome-wide level, novel loci (after excluding literature-based COPD risk loci, **Supplementary Table 6**) identified through TWAS approach included *MROH1* on 8q24.3 and *SYCE1* on 10q26.3. Significant colocalisation was also observed at eight novel loci: *ZDHHC21* on 9p22.3, *CAMK2A* on 5q32, *DMPK* on 19q13.32, *PRR16* on 5q23.1, *MYO15A* on 17p11.2, *TNFRSF10A* on 8p21.3, *BCO1* on 16q23.2, and *HOXC6* on 12q13.13. Finally, four SMR genes were identified including two overlapping with those discovered by colocalisation, namely *CAMK2A* and *DMPK* as well as *BTN3A2* on 6p22.2 and *TRBV30* on 7q34. The biology of these genes and their potential link to COPD pathobiology is provided in **Table 1**, with more details in **Supplementary Table 10**. Interestingly, the top colocalisation (*ZDHHC21*) gene is implicated in lung vascular endothelial barrier integrity.^22^ The p values of the top GWAS SNPs in ICGC located 500 kb up and downstream of genome-wide discovered genes varied from 1.71x10^−5^ to 5.85x10^−4^, suggesting that the largest GWAS on COPD alone was underpowered to identify these genes at genome-wide significance. The findings support the utility of leveraging transcriptome data to uncover biologically relevant genes.

Complementary and as a functional follow-up of GWAS, methods used herein represent an important step to uncover genes whose expression changes in lung tissue are causally related to COPD. In this study, we provided insightful results about causality genes for 60 out of 129 COPD risk loci derived from the literature. For 18 (30%) of these loci, we confirmed the target gene suspected by the GWAS. These include *C1GALT1* on 7p22.1 and *THSD4* on 15q23. The 7p22.1-*C1GALT1* locus was recently reported as one of the 43 new signals for lung function^23^. The 15q23-*THSD4* locus was repeatedly associated with lung function^23,24^ and COPD^10^. In this study, the different integrative methods consistently pointed to these genes as being causally linked to COPD. With the wealth of susceptibility loci being reported (>100 loci), our study provides much-needed information to prioritize follow-up functional studies. In addition, we identified the most likely causal gene for 19 loci (32%) where more than one gene was reported by GWAS. This includes 6p24.3 with two genes suspected from GWAS, namely *BMP6*^23,25^ and *DSP*^10^. Integrative analyses consistently support *DSP* as the causal gene and demonstrate again how our study is narrowing down the investigational space underneath GWAS loci. Finally, for 23 loci (38%) the gene showing the most convincing evidence of causality was not reported by GWAS. As illustrated in **Supplementary Fig. 4**, the causal genes supported in this study were not necessarily the nearest annotated gene to the lead GWAS SNP.

Stringent significant thresholds are used in GWAS to control for false-discovery rate. Many true-positives loci are likely to exist below this threshold. These borderline loci constitute the “grey zone” of GWAS. In this study, we leverage the lung eQTL dataset to uncover true-positives in the grey zone of the largest GWAS on COPD. A total of 20 loci in the grey zone which did not overlap with literature-based COPD risk loci were reported. Interestingly, eSNP-regulated genes (P_eQTL_<0.001) were identified in 3 of these loci including a pseudogene (*LOC100129534*) on 1p36.32, *TM9SF4* (transmembrane 9 superfamily member 4) on 20q11.21 known as an hypoxia-responsive molecule involved in cell adhesion,^26^ and *DNAL1* (dynein axonemal light chain 1) on 14q24.3 implicated in the motility of cilia and with a known homozygous mutation causing primary ciliary dyskinesia characterized by chronic destructive airway disease.^27^ Colocalisation and SMR further supported causality of the pseudogene on 1p36.32, which is classified as a non-coding RNA with unknown function. TWAS genes were also found in liberal loci including *C2CD3* (C2 calcium dependent domain containing 3) on 11q13.4 known to play a role in ciliopathies,^28^ *NUMB* (NUMB, endocytic adaptor protein) on 14q24.3 involved in development and cancer,^29^ and *NOL4L* (nucleolar protein 4 like) on 20q11.21 whose function is largely unknown, but previously implicated in leukemia^30^ and zebrafish development.^31^

For many COPD risk loci, the most likely causal genes were not identified with the current data and will require further investigation. This is consistent with the lower than expected number of variants that colocalized between GWAS on glucose-and insulin-related traits and eQTL in human pancreatic islets and 44 different tissues in the GTEx Portal.^32^ In many cases, the biological mechanisms underlying GWAS loci will not be mediated by eQTL. In this study, we used the most disease-relevant tissue to study COPD, but unavoidably we may have missed gene regulation processes that are specific to other tissues. Studying the whole lung transcriptome of patients undergoing surgery, does not allow to find eQTL specific to a disease-relevant cell type or eQTL that are context dependent, i.e. observed at certain stages of life or disease. It must be emphasized that the results from integrative approaches will require experimental work to confirm the role of identified genes in COPD. In addition, for some COPD risk loci, our results implicate more than one causality gene and further research is needed to explore multiple causal genes in a single locus.

In this study, we leveraged the largest lung eQTL dataset and GWAS on COPD available to provide insights about causality genes and find 12 genes outside GWAS loci as causally related to COPD. By synthesizing the GWAS literature on COPD, we collated 129 non-overlapping risk loci and provided insightful results about the causal gene(s) for 60 of them. Many of these genes were not the closest to, or harboring the lead GWAS variant. We also mapped causality genes in four loci located in the grey zone of GWAS. Overall, by identifying plausible causal COPD genes, this study translates genetic associations into knowledge that is one step closer to clinical applications.

## Acknowledgements

The authors would like to thank the staff at the Quebec Respiratory Health Network Tissue Bank for their valuable assistance with the lung eQTL dataset at Laval University. Ma’en Obeidat is a Fellow of the Parker B. Francis Foundation. He is also a recipient of British Columbia Lung Association Research Grant. Maxime Lamontagne is the recipient of a doctoral studentship from the Fonds de recherche Québec - Santé (FRQS). Jean-Christophe Bérubé is the recipient of doctoral scholarships from the Canadian Respiratory Research Network (CRRN) and FRQS. Ke Hao is partially supported by the National Natural Science Foundation of China (Grant No. 21477087, 91643201) and by the Ministry of Science and Technology of China (Grant No. 2016YFC0206507). Yohan Bossé holds a Canada Research Chair in Genomics of Heart and Lung Diseases. This study was supported by grants from the Chaire de pneumologie de la Fondation JD Bégin de l’Université Laval, the Fondation de l’Institut universitaire de cardiologie et de pneumologie de Québec, the Quebec Respiratory Health Network, and the Canadian Institutes of Health Research (MOP – 123369) to Yohan Bossé. Research reported in this publication was supported by NHLBI R01 HL113264 (M.H.C). The content is solely the responsibility of the authors and does not necessarily represent the official views of the National Institutes of Health. The funders had no role in study design, data collection and analysis, decision to publish, or preparation of the manuscript.

## Author contributions

MO, ML, JCB, DDS, PDP, and YB conceived and designed the study. MHC, BDH, KdeJ, and HMB led the GWAS data collection and analysis of the International COPD Genetics Consortium. DN, KH, WT, MvdB, PJ, ML, DDS, PDP, and YB led the lung eQTL data collection. MO, ML, JCB, and YB performed data analyses. MO, ML, JCB, and YB wrote the manuscript. All authors discussed the results and implications and commented on the manuscript at all stages.

## Competing financial interests

None declared.

## ONLINE METHODS

### Published GWAS risk loci for COPD

**Supplementary Table 4** shows the COPD susceptibility loci identified by review of the literature. This table is an extension of our previous review on the genetics of COPD^4^ and was manually curated by reviewing the published GWAS on COPD, lung function, lung function decline, emphysema, chronic bronchitis, and other-related phenotypes published before March 1, 2017. For each study, we provide the reference, sample size, specific phenotype, suspected susceptibility genes, and key SNPs as reported in the publications. Susceptibility loci were further validated and complemented using the GWAS Catalogue.^33^ Results of GWAS in **Supplementary Table 4** were then arranged by locus in **Supplementary Table 6**. These loci are considered *literature-based COPD risk loci*. In addition, all SNPs with nominal P_GWAS_<1x10^−5^ in ICGC (but sub-genome-wide significant) were considered. These loci are henceforth referred to as *liberal COPD risk loci* and are indicated in **Supplementary Table 8**. For both literature-based and liberal COPD susceptibility loci, the boundaries of each locus were defined as follows: For literature-based COPD loci, key SNPs derived from scientific publications were tabulated for each locus and the locations of the most 5’ and 3’ SNPs were identified. The boundaries of each locus were then defined by adding 500 kb downstream of the most 5’ SNP and 500 kb upstream of the most 3’ SNP. For liberal COPD loci, the boundaries of each locus were defined by a 500 kb region 5’ and 3’ of COPD-associated SNPs with P_GWAS_<1x10^−5^ (i.e. 1 Mb window). When windows overlapped, the intervals were amalgamated into a single interval with 500 kb on either side of each hit. The final boundaries are provided in **Supplementary Tables 6** and **8**.

### ICGC GWAS on COPD

The International COPD Genetics Consortium (ICGC) has recently reported the world’s largest GWAS on COPD risk.^10^ For the current study, only GWAS results for individuals of European ancestry were considered consisting of 20 studies with 13,710 COPD cases and 38,062 controls. Cases were defined by moderate-to-severe airflow limitation based on pre-bronchodilator spirometry measurements (% predicted FEV1<80% and FEV1/FVC<0.7) and GOLD recommendations^1^ indicative of COPD GOLD stage 2 or worse. Genome-wide genotyping data were obtained for cases and controls, and additional SNPs were imputed using the 1000 Genomes reference set. Quality control details have been published previously^10^ and SNPs were considered in the GWAS analysis if they were included in at least 13 of the studies. The GWAS was performed using logistic regression of genotype dosage on COPD case-control status in each cohort separately adjusting for age, sex, smoking status (ever smoking and current smoking), pack-years smoking, and ancestry-based principal components as needed. Results were then meta-analyzed in metal^34^ using fixed effect model with inverse variance weighting. Summary statistics were available for 6,948,071 SNPs including chromosome position, alleles, allele frequencies, p values, effect estimates, and standard errors for all studies evaluated. For ICGC, each cohort obtained approval from appropriate ethical/regulatory bodies; informed consent was obtained for all individuals.

### Lung eQTL mapping study

The lung eQTL study has been previously described.^14-16^ Human lung tissues from subjects who underwent lung surgery were obtained at three academic sites, Laval University, University of British Columbia (UBC), and University of Groningen. All patients provided written informed consent and the study was approved by the ethics committees of the Institut universitaire de cardiologie et de pneumologie de Québec (IUCPQ) and the UBC-Providence Health Care Research Institute Ethics Board for Laval and UBC, respectively. The study protocol was consistent with the Research Code of the University Medical Center Groningen and Dutch national ethical and professional guidelines (‘‘Code of conduct; Dutch federation of biomedical scientific societies’’; http://www.federa.org). Genotyping, gene expression, and lung *cis*-eQTL analyses are described in the online supplementary materials.

### Methods of GWAS-eQTL integration

#### Transcriptome-wide association study (TWAS)

The TWAS was performed using FUSION^21^. The 1,038 individuals for whom both gene expression and genetic variants were measured (i.e. the lung eQTL dataset) were combined with summary-level GWAS data from ICGC to estimate association statistics between gene expression and COPD. Briefly, this method can be conceptualized as having imputed expression data for all cases and controls in ICGC (using the part of expression that can be explained by SNPs in the lung eQTL dataset) and then test for association between imputed gene expression and COPD. To do so, normalized gene expression from Laval, UBC, and Groningen were first combined using ComBat adjustment method^35^ to correct for study site. Second, the genetic values of expression were computed one probe set at a time using SNP genotyping data located 500 kb on both sides of the probe sets using prediction models implemented in FUSION including 1) the single most significant lung eQTL-SNP as the predictor (top1), 2) LASSO regression, and 3) elastic net regression (enet). All probe sets that passed QC in the lung eQTL were evaluated (n=40,359) and 12,474 of them showed significant *cis*-heritability (i.e. part of expression variability that can be explained by SNPs). The expression weights of these *cis*-heritable probe sets were then combined with summary statistics from ICGC to obtain Z-score for each probe set. Genome-wide significant TWAS genes were considered at P_TWAS_<0.0001. A higher cutoff threshold of P_TWAS_<0.05 was used for literature-based and liberal COPD risk loci as we aim to identify the most likely causal gene in these previously established COPD risk loci.

#### Bayesian colocalisation

Summary statistics, more specifically regression coefficients and their variance, from the ICGC GWAS and lung eQTL results were combined using COLOC package version 2.3-6 in R.^19^ Briefly, this method assesses whether two association signals, in this case GWAS and eQTL, are consistent with shared causal variant(s). Default prior probabilities of the software were used, i.e. *p*_*1*_=1x10^−04^, *p*_*2*_=1x10^−04^, *p*_*12*_=1x10^−05^. Genes that demonstrated a high posterior probability (PP4 >75%) indicating that the COPD GWAS and lung eQTL signals colocalized were reported. PP4 > 60% was also considered within literature-based and liberal COPD risk loci.

#### Summary data-based Mendelian randomization (SMR)

GWAS and lung eQTL data were also integrated using the SMR method.^20^ Conceptually this approach is similar to standard Mendelian randomization analysis, where measured variations in genes are used as instrumental variables to test for causative effect of an exposure on disease. Here, the SNPs (instrumental variables) are used to test for the causative effect of gene expression (exposure) on COPD (disease). By default, SMR only considered probe sets with at least one *cis*-eQTL P_eQTL_<5x10^−8^. In this analysis, 8,679 probe sets were evaluated. A cutoff threshold of P_SMR_<0.001 with no evidence of heterogeneity (P_HEIDI_>0.05) was used for genome-wide analysis. For literature-based and liberal COPD risk loci, SMR genes were those with a P_SMR_<0.05 with no evidence of heterogeneity (P_HEIDI_>0.05).

#### eQTL analysis with GWAS SNPs (eSNP)

This method was performed at the individual SNP level and tested whether GWAS SNPs (from literature or liberal COPD risk loci from ICGC) act as lung eQTL. Lung eQTL-regulated genes by GWAS SNPs were considered eSNP-regulated genes. eSNP-regulated genes with P_eQTL_<1x10^−8^ were considered statistically significant. However, eSNP-regulated genes with P_eQTL_<0.05 were also explored as these analyses were performed at previously established COPD loci.

### Data availability

Gene expression data for the lung eQTL dataset are available in the Gene Expression Omnibus repository through accession number GSE23546. The genome-wide association summary statistics from the International COPD Genetics Consortium (ICGC) are available at the database of Genotypes and Phenotypes (dbGaP) under accession phs000179.v5.p2.

